# Albumin-binding dendrimer-conjugated siRNA enables safe and effective gene silencing throughout the central nervous system

**DOI:** 10.64898/2025.12.16.694641

**Authors:** Hassan H. Fakih, Masahiro Ohara, Ashley Summers, Samantha L. Sarli, Karen Kelly, Rosemary Gagnon, Bruktawit Maru, Brianna Bramato, Anastasia Khvorova, Jonathan K. Watts

**Affiliations:** RNA Therapeutics Institute, University of Massachusetts Chan Medical School, Worcester, MA 01605, USA; Boston Children’s Hospital, Harvard University, Boston, MA 02115, USA

## Abstract

Improving siRNA delivery to the central nervous system (CNS) is a major focus for treating the numerous debilitating neurological conditions which have a genetic basis. Here, we present an albumin-binding siRNA based on an amphiphilic dendrimer conjugate (D-siRNA). We demonstrate that D-siRNA achieves effective and homogeneous delivery throughout the CNS following administration into the cerebrospinal fluid (CSF). In mice, a single CSF administration of D-siRNA resulted in potent and durable gene silencing across various brain regions, with effects lasting six months without detectable toxicity. We validate its utility in larger rodents (rats) using intrathecal administration—a clinically relevant route—showing effective and broad delivery and robust silencing. Benchmarking against other clinically relevant siRNA delivery scaffolds revealed that D-siRNA provides comparable delivery and efficacy, with more efficient conversion of gross uptake to functional uptake. These findings support the use of albumin-binding conjugates for brain delivery, and position D-siRNA as a safe, effective, and durable platform for gene silencing in the CNS.

## INTRODUCTION

Oligonucleotide based therapeutics, such as small interfering RNA (siRNA), antisense oligonucleotides (ASO) and splice-switching oligonucleotides (SSO), have been established as a productive therapeutic modality (1-3). Of particular interest for gene silencing are siRNAs, which offer long-lasting durability, with some single doses providing an effect for up to a year, leading to eight approved drugs. However, the currently approved siRNA therapies are limited to the liver, enabled by the robust hepatocyte delivery enabled by the N-acetylgalactosamine (GalNAc) conjugate (2). Current research efforts are intensely focused on expanding the therapeutic potential of oligotherapeutics to extrahepatic tissues, such as muscle, heart, and the central nervous system (CNS) (4). The CNS has drawn sharp focus because many severe neurological diseases have well-characterized pathology stemming from identifiable target genes that oligotherapeutics are ideally positioned to address (5,6). The CNS can be reached by intrathecal administration of oligonucleotides into the cerebrospinal fluid (CSF) which allows for broad distribution throughout brain regions as is necessary for many neurological and neurodegenerative diseases (6).

When ASOs and siRNAs are administered into the CSF, they are quickly cleared due to their small size and hydrophilicity, limiting their cellular uptake and deep brain region penetration (7,8). Strategies to reduce clearance and improve wide distribution and uptake have mainly focused on increasing lipophilicity or cell-surface binding through chemical modification of the siRNA or lipophilic conjugation (7-10). However, lipophilic conjugation is a double-edged sword: While it can minimize clearance and improve cellular uptake (7), highly hydrophobic conjugates can limit diffusion from the injection site and most importantly can induce toxicity (11,12).

Recently, we have reported on the development of an amphiphilic dendrimer conjugate for extrahepatic siRNA delivery (D-siRNA)(12-16). The conjugate’s chemical structure—featuring lipid moieties separated by phosphate groups—imparts its essential amphiphilic nature (13,15). This design successfully balances the need for improved lipophilicity to enhance siRNA circulation and cellular uptake with the need to avoid the toxicity frequently observed with highly lipophilic conjugates, driving effective siRNA activity *in vivo* (12,15,16). Beyond its amphiphilic nature, the delivery benefits of D-siRNA may stem from its capacity for avid and selective albumin binding (15). This feature is particularly significant for CNS distribution, as albumin has been shown to distribute within the brain via the glymphatic system (17), which has been shown to mediate the movement of oligonucleotide therapeutics throughout the brain and spinal cord (18,19). We hypothesized that D-siRNA would achieve effective CNS distribution and transport following CSF administration, with improved distribution to various brain regions and low toxicity. Encouragingly, an independent study by Sorets *et. al*. also recently reported that another albumin-binding ligand gave siRNAs a CNS distribution profile consistent with glymphatic system transport (20).

In this work, we explore and characterize the distribution, efficacy, and safety of the D-siRNA conjugate for CNS delivery. Specifically, we assessed D-siRNA following both intracerebroventricular (ICV) injection and the more clinically relevant intrathecal (IT) route. When benchmarked against the clinically relevant divalent siRNA technology (Di-siRNA)(8,21,22), D-siRNA demonstrated comparable distribution, efficacy, and safety. Following ICV injection, D-siRNA activity was long-lasting (up to six months) and showed potency against multiple targets. In rats, IT injection resulted in wide and effective distribution throughout the CNS, exhibiting no observed toxicity and efficacy lasting at least one month, with trends showing improved potency (lower IC50) compared to Di-siRNA. Fluorescent imaging confirmed colocalization with blood vessels, consistent with glymphatic system transport. This work establishes the amphiphilic-conjugated D-siRNA platform that enables effective and safe oligotherapeutic delivery throughout the CNS.

## MATERIALS AND METHODS

### Synthesis and purification of oligonucleotides

Compounds were synthesized using a MerMade 12 (LGC Biosearch Technologies) synthesizer following standard protocols at 10–20 μmol scales. Standard RNA 2′-O-methyl, 2′-fluoro modifications were used to enhance siRNA stability (ChemGenes and Hongene Biotech). Di-branched sense strands were synthesized on custom-synthesized branchpoint-functionalized controlled pore glass (CPG) as previously described (8), and unconjugated and dendrimer sense strands were synthesized on CPG functionalized with UnyLinker (ChemGenes) as previously reported (15). For the dendrimer sense strand, commercially available amidites (i.e. C6, C12 and symmetrical branching amidite from ChemGenes and Glen Research) were used to build the dendritic moiety on the 5′-end as previously described. All sense strands had a dT2 spacer between the oligonucleotide and the conjugate. Antisense strands were synthesized on CPG functionalized with a Unylinker (ChemGenes), and custom 5′-(*E*)-vinylphosphonate 2′-OMe-uridine CED phosphoramidite (Hongene) was applied to introduce 5′-(*E*)-vinylphosphonate.

All strands were cleaved and deprotected using 28% aqueous ammonium hydroxide solution for 20 hours at 55°C, followed by drying under vacuum at 40°C, and resuspension in Millipore H_2_O. Oligonucleotides were purified using an Agilent 1290 Infinity II HPLC (Agilent Technologies) on a C18 column for lipid-conjugated (dendrimer) sense strands and an ion-exchange column for the other strands. Purified oligonucleotides were desalted by size-exclusion chromatography and characterized by LC-MS analysis on an Agilent 6530 accurate-mass quadrupole time-of-flight (Q-TOF) LC/MS (Agilent Technologies). The sequences and modifications of the oligonucleotides are shown in **Supplementary Table S1**. HTT and APP sequences were acquired from Alterman *et al*.*(8)* and Sarli *et al. (12)*, JAK1 from Tang *et al. (23)*, and MECP2 from Hariharan *et al*. (24).

### Duplex formation

Equimolar amounts of antisense and sense strands were prepared in water and incubated at 95°C for 5 minutes, cooled to room temperature and at stored at 4°C if not used immediately. The duplexes were then dried and resuspended in artificial CSF (aCSF, 137 mM NaCl, 5 mM KCl, 20 mM glucose, 8 mM HEPES, 2.3 mM CaCl_2_, 1.3 mM MgCl_2_, pH 7.4 – for mice aCSF was supplemented up to 14 mM CaCl_2_, 2 mM MgCl_2_ (25)). Duplexes were resuspended to a concentration of 10 nmol/5μL, 10 nmol/10μL or 80nmol/70μL, depending on the individual experiment. To validate efficient duplex formation, 20 pmol of duplex was loaded onto a non-denaturing 20% tris-borate-EDTA (TBE) gel (Invitrogen # EC63155BOX) and run at constant 180V for 1 hour. The gel was then washed with deionized water for 10 minutes, stained with 1X SYBR Gold Nucleic Acid Gel Stain (Invitrogen # S11494) for 10 minutes or 1X SYBR Safe Nucleic Acid Gel Stain (Invitrogen # S33102) for 15 minutes, and washed again. Bands were visualized on the ChemiDoc MP Imaging system (Bio-Rad # 17001402).

### Animal experiments

All animal procedures were conducted according to the Institutional Animal Care and Use Committee (IACUC) protocols of the University of Massachusetts Chan Medical School (IACUC protocols 202000010 and 202100196) and in accordance with the National Institutes of Health Guide for the Care and Use of Laboratory Animals. Animals were housed and maintained in pathogen-free animal facilities at UMass Chan Medical School with a 12-h light/12-h dark cycle at a controlled temperature (23 ± 1°C), standard humidity (50% ± 20%), and free access to food and water.

### Stereotactic intracerebroventricular (ICV) injections in mice

ICV injections were performed as previously described(26). Briefly, C57BL/6NJ male mice (Jackson Laboratory, Strain # 005304) aged 10-12 weeks were anesthetized with 2.5% isoflurane in oxygen. Hair at surgical sites was shaved with an electric razor and a sterile surgical field was set up around the animal’s head. A burr was used to drill a small hole at the following coordinates relative to bregma: −0.2 mm posterior, 1.0 mm mediolaterally. The needle was then placed at a depth of −2.5 mm ventrally. A volume of 5 μL was injected per ventricle at a rate of 750 nl/min. Following injection, mice were monitored until sternal. All doses are based on molar amounts of active siRNA (antisense strand)

### Intrathecal delivery of oligonucleotides to rats

Twelve-week-old Sprague–Dawley rats (Charles River Laboratories, Strain # 001) were used for the experiments. Intrathecal administration was performed with slight modifications from previous reports (18). Briefly, after isoflurane anesthesia, buprenorphine (0.1 mg/kg) was administered subcutaneously. Using a 50-mL conical tube, the rat was placed in the prone position with the spine flexed, and a skin incision was made. An incision was then made through the visible muscle layer, and the L6 lumbar vertebra was identified. A guide cannula (SAI Infusion Technologies) with an inserted 23-gauge needle (Becton Dickinson) was advanced anterior to the L6 vertebra and positioned in place. Subsequently, a catheter–wire assembly (SAI Infusion Technologies) was inserted through the guide cannula into the spinal column. The guide cannula and stylet wire were removed sequentially, leaving only the catheter in the spinal column. The catheter was gently pulled back until the black mark located 2 cm from the tip aligned with the spinal column. Thereafter, 70 μL of aCSF or siRNA was administered through the catheter tip over a period of more than 30 seconds. All surgical procedures were performed under sterile conditions. All doses are based on molar amounts of active siRNA (antisense strand)

### Limb paralysis scoring after IT injection

Functional abnormalities following IT administration have been reported with oligonucleotide administration (27). We assessed limb paralysis scoring 24 hours (Figure 3) or 24 hours and 6 days (Figure S4) after intrathecal injection of the Htt-siRNAs using a previously reported scoring system to evaluate potential motor phenotypes (28). The examiner was blinded to the treatment allocation.

**Figure 1.**
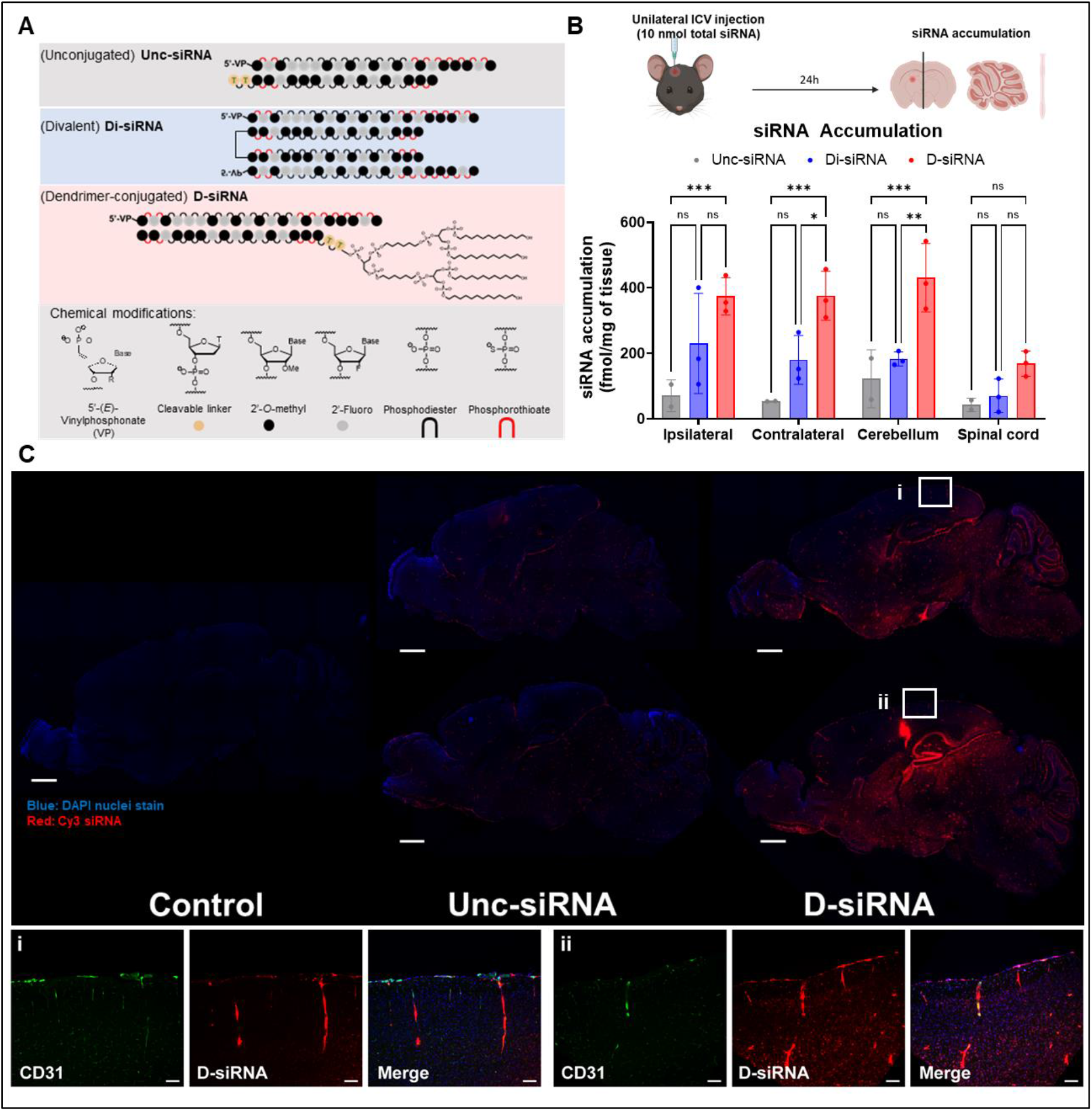
Dendrimer-conjugated D-siRNA achieves widespread delivery in the CNS following ICV injection. **(A)** Schematic of various siRNA scaffolds (unconjugated, divalent and dendrimer-conjugated) assessed for brain distribution, with the chemical modifications used to stabilize the siRNA. **(B)** The siRNA scaffolds were injected unilaterally in the lateral ventricle, followed by collection of various central nervous system (CNS) tissues and siRNA quantification. **(C) (top)** Representative fluorescence imaging of brain tissue 48h post-ICV injection of Cy3 labeled Unc- or D-siRNAs (red=Cy3 siRNA, blue= DAPI staining). **(bottom)** Colocalization of D-siRNA with vascular marker CD31 (green). All study groups are n=3 mice/group. Statistical analysis was performed in GraphPad Prism: two-way ANOVA analysis followed by Tukey’s multiple comparisons across all groups (ns = non-significant, * p<0.05, ** p<0.01, *** p<0.001, **** p<0.0001. (Data represented as mean ± S.D.; tissue accumulation measured by PNA assay; scale bars represent 1 mm in the top panel of **(C)** and 100 µm in the bottom panel of **(C)**; graphical elements created with Biorender).

**Figure 2.**
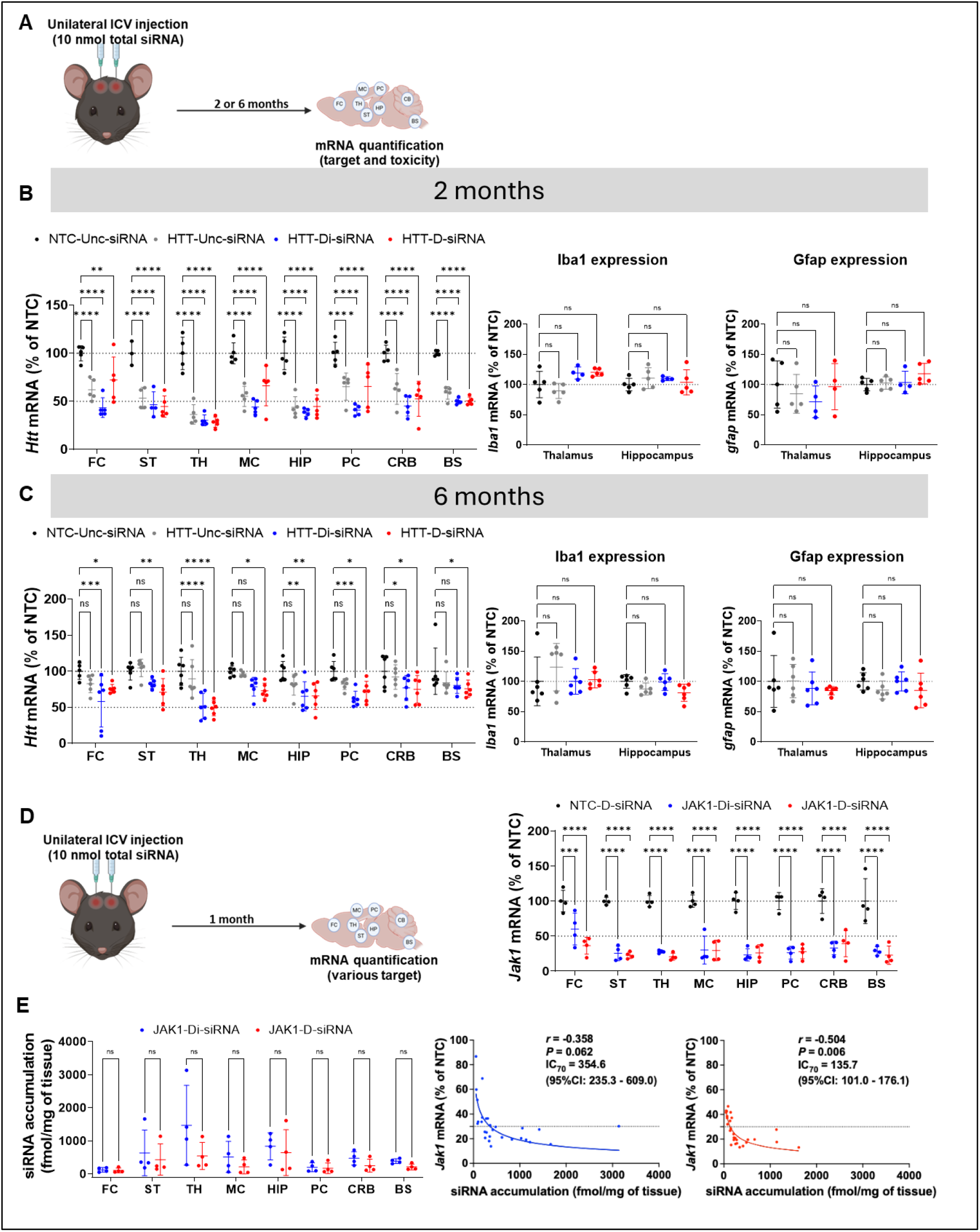
D-siRNA achieves efficacy in multiple brain regions with effects lasting six months after a single injection with no observed toxicity. **(A)** Schematic of *Htt*-targeting siRNA efficacy study following bilateral ICV injection of 10 nmol dose at two- or six-months post injection. Data at **(B)** two-months and **(C)** six-months post injection showing silencing of *Htt* mRNA in various brain regions (left) and levels of neurotoxicity markers mRNA (*Iba1* for microgliosis and *gfap* for astrogliosis. **(D)** Silencing data of *Jak1* mRNA following bilateral ICV injection of 10 nmol dose at one-month post injection, and **(E)** siRNA accumulation comparing D-siRNA and Di-siRNA (left) with PK/PD correlation (right). All study groups are n=4-6 mice/group. Statistical analysis was performed in GraphPad Prism: two-way ANOVA analysis followed by Dunnet’s multiple comparisons of all groups against NTC control group (ns = non-significant, * p<0.05, ** p<0.01, *** p<0.001, **** p<0.0001). (Data represented as mean ± S.D.; mRNA measured by the Quantigene 2.0 Assay; tissue accumulation measured by PNA assay; dotted line on all graphs represents 100% and 50% remaining mRNA protein expression, and for correlation figures represents 30% remaining expression; graphical elements created with Biorender).

**Figure 3.**
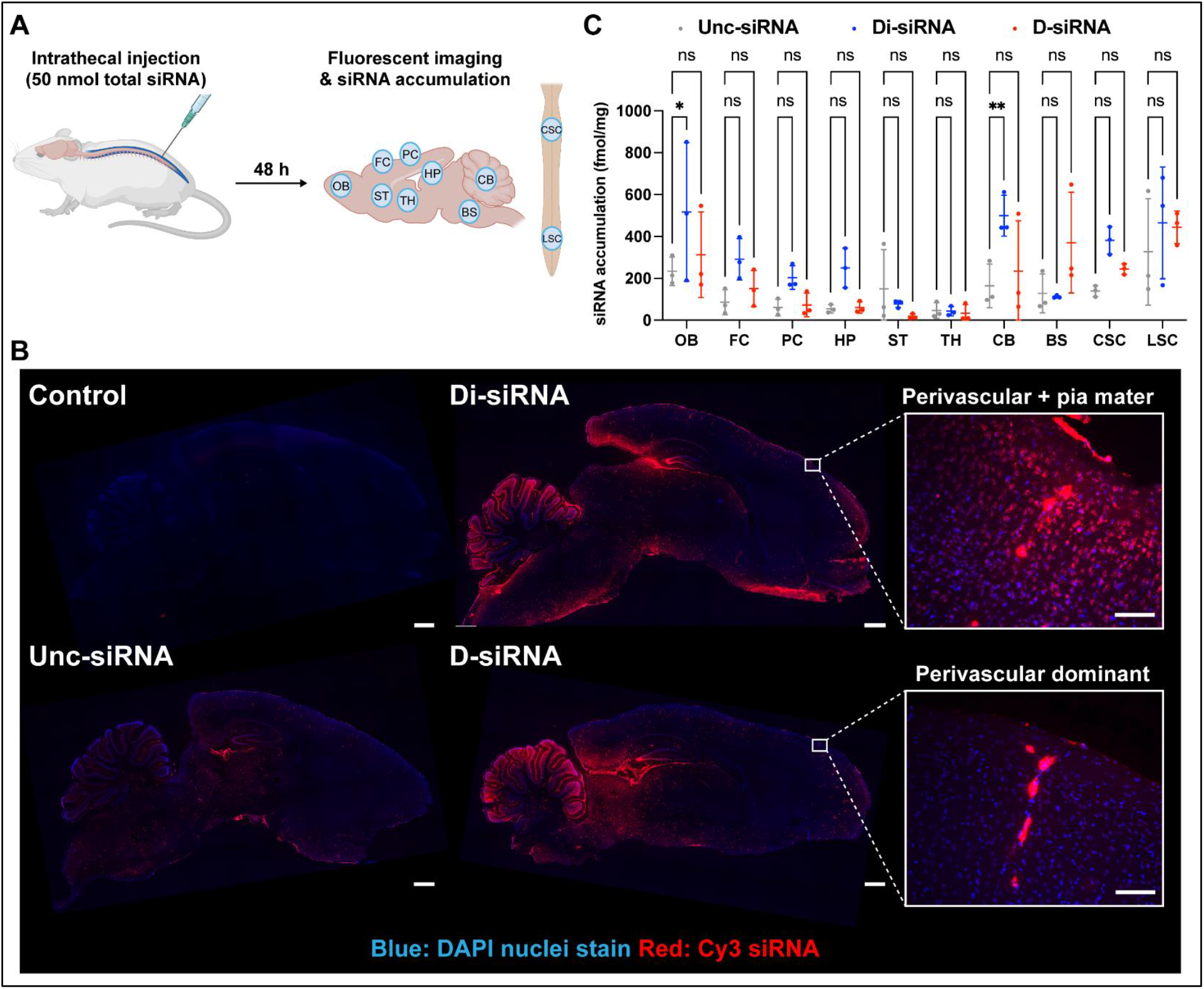
D-siRNA enables safe and widespread distribution via intrathecal injection in rats. **(A)** Experimental schematic for the study of intrathecal injection of 50 nmol dose of Cy3-labeled siRNAs; accumulation and distribution were assessed 48 hours after IT injection of Unc-, Di-, and D-siRNA. **(B)** Representative brain images 48 hours following IT injection. **(C)** siRNA accumulation across the CNS regions. Statistical analysis was performed in GraphPad Prism: two-way ANOVA analysis followed by Dunnet’s multiple comparisons of all groups against UNC group (ns = non-significant, * p<0.05, ** p<0.01, *** p<0.001, **** p<0.0001). (Data represented as mean ± S.D.; tissue accumulation measured by PNA assay, GFAP quantification by immunostaining, limb paralysis scoring was blinded; scale bars represent 1 mm **(C)** and 100 µm in the inset in **(C);** graphical elements created with Biorender).

### mRNA quantification and tissue processing

mRNA expression was quantified with the QuantiGene Singleplex Assay Kit (Invitrogen # QS0016), a hybridization-based assay, as previously described (29,30). Cultured cells were lysed in a total volume of 250 µL diluted lysis mixture consisting of one part lysis mixture, two parts deionized H2O and 0.167 mg/mL proteinase K (Invitrogen # AM2548). Tissue lysates were thoroughly mixed by pipetting up and down 15 - 20 times. Probe set validation was performed for each tissue to determine an appropriate lysate volume that would allow unambiguous and linear mRNA detection. The corresponding volumes were transferred to the capture plate as indicated in the manufacturer’s protocol (Invitrogen # MAN0018628), and the protocol was followed further. Luminescence detection was performed using the SpectroMax M5 Microplate Reader (Avantor).

For mouse tissue: At sacrifice, the brain was harvested and placed in a brain matrix (Braintree Scientific). A 1-mm slice was then taken from the regions of interest and 1.5 × 1.5-mm punches were taken and placed in RNALater (935 ml MilliQ water, 700 g ammonium sulfate, 25 ml 1M Sodium Citrate, 40 ml 0.5 M EDTA, pH 5.2 using H_2_SO_4_) for 24 h at 4°C, then frozen at −80°C until processing for subsequent experiments. Punch biopsies were dried gently, weighed and added to 500 μL of homogenizing solution (Invitrogen; QS0517) containing 0.2 mg/mL proteinase K (Invitrogen, #AM3546), in a QIAGEN collection microtube holding a 3-mm tungsten bead. The tissues were then homogenized for 10 min under 25 Hz of frequency using a QIAGEN TissueLyser II, followed by incubating at 55 °C for 30 min. Lysate was then used as described above for mRNA quantification, or for siRNA accumulation (described below) or stored at -20 °C. The following QuantiGene Singleplex RNA probes were used from ThermoFisher Scientific: Htt (Murine, SB-14150), Hprt (Murine, SB-15463), Jak1 (Murine, SB-3029714), Mecp2 (Murine, SB-11697) Gfap (Murine, SB-14051), Iba-1 (Murine, SB-3027744).

For rat tissue: Rats were euthanized at the specified time point post-injection by CO_2_ inhalation; cardiac perfusion with ice-cold PBS was performed, followed by removal of the brain, spinal cord, liver, and kidney. Brains were oriented in an adjustable brain matrix (Braintree Scientific), and 2 mm sagittal sections were collected through the regions of interest; 3-mm biopsy punches were collected from the specific brain areas. Tissue punches were stored in RNAlater overnight at 4°C. For the evaluation of gene silencing, tissue punches were suspended in Affymetrix homogenizing solution containing proteinase K and mechanically dissociated using a Qiagen Tissuelyser with a 2-mm tungsten carbide bead. The tubes were then incubated in a water bath at 65 °C until all tubes appeared transparent. Tubes were centrifuged (16,000 xG, 15 minutes) and supernatant transferred to a 96 well plate for storage at -80. The following QuantiGene Singleplex RNA probes were used from ThermoFisher Scientific: Htt (Rat, SC-34753), Hprt (Rat, SC-14898), App (Rat, SC-15981).

For the evaluation of Iba1 and Gfap mRNA levels in rats, lumbar spinal cord samples were homogenized in TRIzol reagent (Invitrogen) using a Qiagen Tissuelyser with a 3 mm tungsten carbide bead. RNA was extracted using RNA Clean & Concentrator Kit (Zymo Research, Cat. #R1016) and quantified on a Nanodrop (ThermoFisher Scientific). For reverse transcription, 1 µg of RNA was used with the High-Capacity cDNA Reverse Transcription kit (Applied Biosystems, Cat. #4368813) per the manufacturer’s protocol. qRT-PCR was carried out in technical duplicates using iTaq Universal SYBR Green Supermix (Bio-Rad, Cat. #1725122) on Bio-Rad CFX-96 real time machine using gene-specific primers from Integrated DNA Technologies (IDT, Coralville, Iowa, USA): Gfap forward primer: 5’ GTTAAGCTAGCCCTGGACATC 3’; Gfap reverse primer: GATCTGGAGGTTGGAGAAAGTC; Iba1 forward primer: 5’ TCCGAGGAGACGTTCAGTTA 3’; Iba1 reverse primer: 5’ GTTGGCTTCTGGTGTTCTTTG 3’; Gapdh forward primer: 5’ ACAAGATGGTGAAGGTCGGTG 3’; Gapdh reverse primer: 5’ ACCATGTAGTTGAGGTCAATGAAGG 3’.

### siRNA tissue accumulation quantification

To quantify siRNA accumulated in tissues following systemic administration, PNA hybridization assay was used on the lysate prepared for mRNA quantification, as previously described with slight modification (29-31). All samples that were quantified were weighed to calculate siRNA accumulation per milligram of tissue. Briefly, the accumulation was quantified using custom Alexa488-labeled fluorescent PNA oligonucleotide probes that are fully complementary to the antisense strand of interest (PNABio). The probes used are Jak1 (Alexa488-OO-CATCAGCTACAAGCGAT), Mecp2 (Alexa488-OO-ATCTGACAAAGCTTCCCGATA) and App (Alexa488-OO-ATCAATTACCAAGAATTCTCA). To hybridize to the antisense strand of target siRNAs, lysates were annealed with the corresponding PNA probe (95 °C for 5 mins, followed by 5 mins at 55 °C and 5 mins at 4 °C). Then, lysates were cleaned by precipitating out Sodium dodecyl sulphate by adding 50 μL of 3 M potassium chloride followed by centrifugation for 15 min at 5000×g. The clear supernatant containing the bound antisense strand was then collected. Anion-exchanged chromatography was used to analyze the sample mixtures on an Agilent 1260 Infinity quad-pump HPLC with a 1260 FLD fluorescent detector, and quantification was based on compared to standard curves of spike-in siRNA in tissue lysate as previously described (4).

### Rat protein quantification with Western blots

Rat brain samples were homogenized and lysed in ice-cold RIPA buffer (Boston BioProducts, Cat. #BP-115S) supplemented with Halt protease and phosphatase inhibitor cocktail (Life Technologies, Cat. #78441). Lysates were centrifuged at 12,000 rpm for 10 min at 4 °C, and the supernatant was collected into fresh microcentrifuge tubes, leaving the pellet behind. Protein concentrations were determined using a BCA protein assay (ThermoFisher Scientific, Cat. #23225). Equal amounts of protein (30 µg per sample) were mixed with 4× Laemmli SDS sample buffer (Thermo Scientific Chemicals, Cat. #J60015.AD) and boiled at 95 °C for 5 min. Samples were vortexed and briefly centrifuged before electrophoresis. Proteins were separated on 4–20% Tris– Glycine SDS–PAGE gels (Bio-Rad, Cat. #4561094) in running buffer (Bio-Rad, Cat. #1610732). Electrophoresis was performed at 50 V for 5 min to allow stacking, followed by 100 V for 1 h. Proteins were transferred to nitrocellulose membranes (Bio-Rad, Cat. #162-0115) using a semi-dry transfer apparatus. Transfer was carried out at 20 V for 1 h at room temperature. Then, membranes were blocked with Intercept Blocking Buffer (LI-COR, Cat. #927-60001) at room temperature for 1 h with agitation, followed by overnight incubation at 4 °C with primary antibodies diluted in blocking buffer (1:1,000 for APP (Cell Signaling, Cat. #2452S); 1:2,000 for GAPDH (Sigma-Aldrich, Cat. #MAB374)). After three washes with 1× TBST (10 min each), membranes were incubated with IRDye 680 anti-rabbit antibody for APP (1:5,000; LICORbio, Cat. #925-68073) and IRDye 800 anti-mouse antibody for GAPDH (1:5,000; LICORbio, Cat. #926-32210) for 1 h at room temperature. Membranes were then washed three additional times with 1× TBST. The bands were visualized using the LI-COR imaging system according to the manufacturer’s instructions.

### Histology

Mice or rats were perfused with ice-cold PBS followed by ice-cold 4% paraformaldehyde (PFA) (Thermo Scientific Chemicals, Cat. #J19943-K2). Brains and spinal cords were post-fixed in 4% PFA at 4 °C overnight, and subsequently cryoprotected in 30% sucrose at 4 °C overnight. For experiments examining brain distribution in rats, brain hemispheres were initially preserved in RNAlater and stored at –80 °C, followed by processing using the same protocol described above. Tissues were embedded in OCT compound (Sakura Finetek USA), and frozen sections were cut at a thickness of 20 µm using a cryostat (Thermo Fisher). Sections were mounted onto glass slides (MATSUNAMI, Cat. # SUMGP11). In experiments to assess siRNA distribution, tissue sections were air-dried at room temperature, washed three times with PBST, and coverslipped using a mounting medium containing DAPI (VECTOR laboratories, Cat. #H-1200-10). For immunofluorescence, Sections were washed three times with 1× TBST and incubated in 5% goat serum in TBST for 60 min at room temperature to block nonspecific binding. Samples were then incubated with anti-CD31 antibody (1:200; Cell Signaling Technology, Cat. #15585T) or anti-GFAP antibody (1:200; Cell Signaling Technology, Cat. #80788S) at 4 °C overnight, followed by three washes with TBST. Subsequently, sections were incubated with Alexa Fluor 488-conjugated anti-rabbit IgG secondary antibody (1:500 for CD31 staining and 1:1000 for GFAP staining; Cell Signaling Technology, Cat. #4412S) diluted in 5% goat serum/TBST for 1.5 h at room temperature. After three additional washes with TBST, sections were mounted using a DAPI-containing mounting medium (VECTOR laboratories, Cat. #H-1200-10). For the quantitative evaluation of GFAP-stained area, we used the Analyze Particle tool in ImageJ as previously described (32). Regions of interest (333 x 333 µm) were placed in the left and right anterior horns of the lumbar spinal cord, and the stained areas were compared.

### Blood diagnostics

Blood sampling for blood diagnostics was performed when mice were terminated at 24h post injection, via cheek bleed. In total, 200 μL of blood was collected in a lithium heparin-coated BD Microtainer tube (BD, #365965) for blood chemistry test. 100 μL of blood was collected in a K2 EDTA-coated BD Microtainer tube (BD, #365974) for a complete blood count (CBC) test. Blood chemistry and CBC diagnostics were conducted by the Diagnostics Laboratory in the Department of Animal Medicine at UMass Chan Medical School.

### Graphs and statistical analyses

Data were analyzed using GraphPad Prism 10.1.2 software for Windows (GraphPad Software, Inc., San Diego, CA). For each independent experiment, the levels mRNA silencing was normalized to the mean of the control NTC group. Data were analyzed using a one-way or two-way ANOVA with post hoc multiple comparisons as specified in the figure captions. IC_50_ and IC_70_ values were obtained from 4-parameter logistic fits (nonlinear regression), with Bottom and Top constrained to 0 and 100, respectively, and the “Find ECanything” function was used to determine the concentrations corresponding to 50% and 70% knockdown on the fitted curves. Asterisks (*) denote gene expression significance (^*^*p* <0.05, ^**^*p* <0.01, ^***^*p* <0.001, ^****^*p* <0.0001). Graphs are plotted as mean ± standard deviation.

## RESULTS

### D-siRNA achieves brain-wide distribution in mice following administration into the CSF

To evaluate the CNS distribution of different siRNA chemistries, mice received a unilateral intracerebroventricular (ICV) injection of 10 nmol of Cy3 labeled, fully chemically modified unconjugated (Unc-siRNA), divalent (Di-siRNA), or dendrimer-conjugated (D-siRNA) siRNAs (Figure 1A). This unilateral injection allowed for the assessment of distribution to both hemispheres and the broader CNS parenchyma (Figure 1B). Twenty-four hours post-injection, the accumulation of siRNA was quantified in the ipsilateral and contralateral hemispheres, cerebellum, and spinal cord. Both Di- and D-siRNA significantly enhanced siRNA distribution compared to Unc-siRNA across all collected tissues (***p<0.001), with the exception of the spinal cord. Distribution levels of Di- and D-siRNA were generally comparable throughout the CNS, though D-siRNA showed an improvement in the contralateral hemisphere and cerebellum compared to Di-siRNA (400 vs. 200 fmol/mg *p< 0.05 and 400 vs 200 fmol/mg **p<0.01, respectively). We also assessed if such administration has any impact on complete blood count or blood chemistry as readouts of any potential systemic toxicity and found no alterations to any of the markers measured compared to controls (Supplementary Figure 1).

Fluorescence imaging of brain sections further confirmed the dendrimer conjugate’s ability to improve both the distribution and retention of the siRNA cargo relative to Unc-siRNA (Figure 1C). When co-staining for CD31 (blood vessels), a strong correlation was observed with the Cy3-labeled D-siRNA. Collectively, these data demonstrate that the albumin-binding D-siRNA achieves wide CNS distribution, comparable to that of clinical-stage divalent siRNA chemistry, and representing a significant improvement relative to unconjugated siRNAs.

### D-siRNA supports safe, potent and durable activity in the CNS

Following the distribution assessment, we evaluated the efficacy and safety of D-siRNA in the mouse CNS. Mice were bilaterally injected (ICV) with a total dose of 10 nmol (5 nmol/ventricle) of a non-targeting control (NTC) or previously developed Huntingtin (*Htt)-*targeting sequence (8,12), comparing Unc-, Di-, or D-siRNA chemistries (Figure 2A). Efficacy was assessed by quantifying *Htt* mRNA expression, while toxicity was monitored using *Iba1* (microglia activation) and *Gfap* (astrogliosis) mRNA expression (8). This was done at both at two months (Figure 2B) and six months post injection (Figure 2B), to evaluate durability.

At two months post-injection (Figure 2B), all chemistries showed similarly robust silencing of *Htt* mRNA compared to the NTC control (Figure 2B, left panel). Analysis of toxicity indicators *iba1 and gfap* in the thalamus and hippocampus—regions closest to the injection site with the highest siRNA accumulation and activity (8)— showed no observed changes, supporting the safety of D-siRNA (Figure 2B, right panel).

By six months post injection (Figure 2C), Unc-siRNA lost its activity in all brain regions, whereas both Di- and D-siRNA maintained significant silencing in most regions. In striatum, medial cortex and brain stem, only the D-siRNA maintained significant *Htt* mRNA silencing at six months (∼25%, *p<0.05 for all regions, compared to NTC) (Figure 2C, left panel). Toxicity markers remained unchanged across all the groups, further supporting the long-safety of D-siRNA with a profile similar to that of Di-siRNA. Overall, the data demonstrated comparable silencing durability and safety of D-siRNA to that of Di-siRNA, with durability enhancement compared to Unc-siRNA which lost efficacy at six months post injection.

To confirm the broad applicability of the D-siRNA platform, the dendrimer was conjugated to siRNAs targeting *Jak1* (Figure 2D) and *Mecp2* (Supplementary Figure 2), both using previously identified cross-species-reactive sequences (23,24). Following a bilateral ICV injection (10 nmol total dose), the activity of D-siRNA was compared to Di-siRNA one-month post-injection (Figure 2D, left panel). Both constructs demonstrated robust silencing efficacy, achieving approximately ∼75% *Jak1* mRNA silencing across all regions (Figure 2D, right).

Assessment of JAK1-siRNA accumulation showed no significant difference in accumulation between D-siRNA and Di-siRNA across various brain regions (Figure 2E, left panel). Plotting the *Jak1* mRNA silencing versus siRNA accumulation allowed for a direct comparison of the PK/PD relationship between the two chemistries (Figure 2E, right panel). Both constructs showed a strong PK/PD correlation, but D-siRNA achieved a significantly lower IC70 (135.7 fmol/mg with 95% CI 101–176.1) compared to Di-siRNA (354.6 fmol/mg with 95% CI 235.3–609.0), indicating enhanced delivery efficiency (i.e. conversion of gross uptake to functional uptake). Similar results were obtained with *Mecp2*-targeting siRNAs where both chemistries achieved potent efficacy of ∼75% silencing across all regions (Supplementary Figure 2).

### D-siRNA enables safe and brain-wide distribution via intrathecal administration in rats

Based on the promising ICV results in mice, we next evaluated intrathecal (IT) administration, a delivery method more widely used in the clinical setting for oligonucleotides in the CNS (2). Although ICV dosing can achieve efficacy in deep brain regions due to these areas being close to the injection site, IT delivery has been reported to have limited access to deep structures (33,34). Therefore, we assessed whether D-siRNA would retain distribution and efficacy across the CNS following IT administration. To first examine CNS distribution, we administered Cy3-labeled siRNAs intrathecally to rats (50 nmol, Unc-, Di- and D-siRNAs) and evaluated their distribution in the brain and spinal cord 48h post injection.

In the brain, the Unc-siRNA showed weaker signal than either Di- or D-siRNA. The Di-siRNA showed a marked signal intensity in brain parenchyma, in addition to exhibiting a bright signal along the pia mater and perivascular structures (Figure 3B). Interestingly, the D-siRNA showed much more intense signal along perivascular structures (Figure 3B), consistent with the hypothesis that the albumin-binding nature of this dendrimeric conjugate facilitates delivery through the glymphatic system. A similar pattern was observed in the lumbar spinal cord: D-siRNA produced signal levels comparable to Di-siRNA and stronger than Unc-siRNA (Figure S3A). Quantification of siRNA accumulation demonstrated that D-siRNA accumulated most strongly in the lumbar spinal cord and, overall, tended to accumulate similarly to Di-siRNA (Figure 3C).

For toxicity assessment, we evaluated GFAP immunofluorescence in the lumbar spinal cord, the region with the highest D-siRNA accumulation, as it has been reported to transiently increase with oligonucleotide administration (8). No obvious differences in GFAP staining area were observed between groups (Figure S3B, C). Furthermore, motor scoring at 24 hours post-injection revealed no detectable functional abnormalities (Figure S3D).

### Robust knockdown of *App* with intrathecally administered D-siRNA

Based on the favorable safety profile and distribution of D-siRNA following IT administration, we then assessed the silencing efficacy using siRNAs targeting a disease-relevant gene. We compared the pharmacodynamics and pharmacokinetics of siRNAs targeting amyloid beta (Aβ) precursor protein gene (*App*), a target for which siRNAs are already in clinical development as a treatment for Alzheimer’s disease (7,35). App-targeting Unc-, Di- and D-siRNAs were IT administered at 80 nmol dose of each compound into rats and evaluated readouts at one month post injection (Figure 4A). D-siRNA showed robust target gene silencing of approximately 50-70% across almost all brain regions, similar to Di-siRNA, and achieved significant knockdown even in deep brain regions where Unc-siRNA showed limited silencing (Figure 4B). At the protein level, D-siRNA achieved a significant reduction of APP throughout the cortex, hippocampus, and brainstem (Figure 4C).

**Figure 4.**
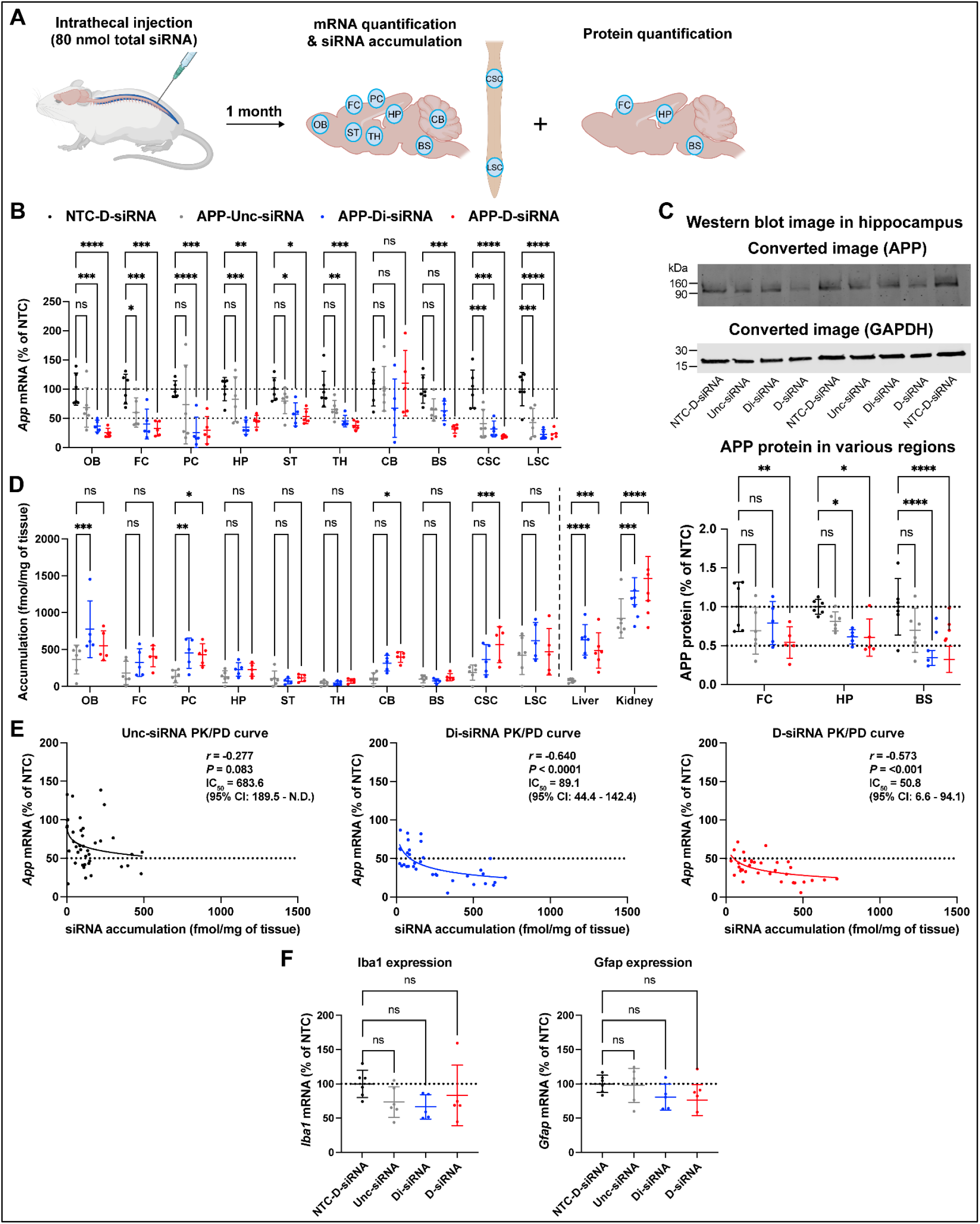
D-siRNA enables efficacious and safe siRNA delivery in rats via intrathecal injection. **(A)** Experimental schematic for the study of intrathecal injection of 80 nmol dose of App-targeting siRNAs; mRNA levels, siRNA accumulation, and protein expression were assessed one month after IT injection of Unc-, Di-, and D-siRNA. **(B)** Silencing data of *App* mRNA in various CNS regions. **(C)** Representative Western blot images **(top)** and quantification of APP protein levels in the frontal cortex, hippocampus, and brainstem **(bottom)**. Uncropped blot images and additional primary Western blot data are included in Supporting Figure S5. **(D)** siRNA accumulation across the CNS regions, liver, and kidney. **(E)** Relationships between siRNA accumulation and App mRNA levels. **(F)** Levels of neurotoxicity mRNA markers in the lumbar spinal cord (*Iba1* for microgliosis and *Gfap* for astrogliosis). All study groups are n=5 or 6 rats/group. Statistical analysis was performed in GraphPad Prism: two-way ANOVA analysis followed by Dunnet’s multiple comparisons of all groups against UNC group (ns = non-significant, * p<0.05, ** p<0.01, *** p<0.001, **** p<0.0001). (Data represented as mean ± S.D.; tissue accumulation measured by PNA assay; graphical elements created with Biorender).

In terms of siRNA accumulation, as expected, both Di-siRNA and D-siRNA showed an overall higher accumulation than Unc-siRNA (****p<0.0001 for Di- and D-siRNA vs Unc-siRNA, in all CNS regions), though the differences were statistically significant only in some individual brain regions (Figure 4D). Interestingly, when PK/PD relationships were evaluated—excluding the cerebellum which shows low silencing with high accumulation as previously reported (36,37)—both D-siRNA and Di-siRNA showed substantially better silencing per tissue accumulation than Unc-siRNA. D-siRNA showed the highest efficiency of converting gross uptake to functional silencing, with an in-tissue IC_50_ of 50.8 fmol/mg tissue; 95% CI: 6.6 – 94.1 (Figure 4E). This result is in broad concordance with the data observed by ICV injection in mouse (Figure 2). No evidence of neuroinflammation was observed based on (*Iba1* and *Gfap)* mRNA levels (Figure 4F).

The broad applicability of the D-siRNA was further confirmed by silencing *Htt* as an additional target. Cy3-labeled, *Htt-*targeting siRNAs administered intrathecally to rat at a relatively low dose (50 nmol) showed moderate but overall significant efficacy at one month post injection (Figure S4A - two-way ANOVA, non-significant for Unc-siRNA, *P* ***p< 0.0001 for Di-siRNA and D-siRNA relative to NTC-siRNA). Importantly, These compounds did not induce any obvious motor phenotypes at 24 hours or 6 days after administration (Figure S4C). Furthermore, neuroinflammation markers (*Iba1* and *Gfap)* showed no changes between groups, even in the lumbar spinal cord where accumulation was the highest (Figure S4D), at one-month post injection.

## DISCUSSION

Oligonucleotide therapeutics show promise to treat neurological diseases in the CNS. However, the CNS is a delicate and sensitive system, and some modifications and conjugates that work for systemic administration show toxicity when delivered to the CNS (25,38,39). Lipophilic conjugation has been shown to improve PK/PD of oligotherapeutics (7,40,41), but highly lipophilic conjugates tend to increase toxicity (seizure-like phenotypes) and limited brain distribution when administered in the brain parenchyma (12,42). Lipophilic conjugates with more moderate hydrophobicity appear to have improved safety profile and wider brain distribution (7,12). Here, we showed that an amphiphilic conjugate that selectively and avidly binds albumin has attractive safety and PK/PD properties for CNS delivery. Another albumin-binding ligand was also recently shown to have promising delivery properties in the CNS (20).

D-siRNAs demonstrated a promising therapeutic profile in the CNS, comparable to the divalent siRNA chemistry which has already received IND approval and characterized by broad distribution and extended durability without compromising safety. The ability of D-siRNA to achieve significant contralateral accumulation following unilateral ICV administration suggests that the conjugate overcomes the diffusion limitations often observed with highly lipophilic modifications (42). We attribute this improved parenchymal permeation to the albumin-binding capacity of the dendrimer, which likely exploits glymphatic transport for wider convective distribution, a mechanism consistent with other albumin-binding constructs (20) and supported in our work by the observed strong colocalization with blood vessels. Notably, the data reveals a divergence between tissue pharmacokinetics (PK) and cellular pharmacodynamics (PD): while dendrimer and divalent chemistries exhibited similar tissue-level accumulation, the dendrimer conjugate achieved a significantly better conversion of gross uptake to functional uptake (lower in-tissue IC_70_). We do not know whether the efficient cellular uptake and endosomal escape shown here are related to the albumin-binding ability of the dendrimer conjugate or simply to its amphiphilic nature (40,43), though the known ability of albumin to undergo transcytosis may be relevant (44).

The translational potential of this platform is further supported by intrathecal (IT) administration in rats, more closely mirroring clinical protocols. The transition to a larger rodent species (∼5-fold brain volume increase) did not diminish efficacy; D-siRNA maintained potent silencing of the Alzheimer’s target *App* across the spinal cord and brain regions. Consistent with ICV findings, IT administration yielded a lower in-tissue IC_50_ compared to divalent siRNAs, reinforcing the hypothesis that the amphiphilic dendrimer improves the efficiency of conversion of gross uptake to functional uptake. Crucially, the absence of neurotoxicity markers (neuroinflammation or motor deficits) across both routes confirms that this enhanced potency does not come at the cost of the safety profile.

As the landscape of CNS oligotherapeutics evolves, there is a distinct drive toward modalities that enable effective brain delivery via systemic administration. The most advanced of these technologies are transferrin receptor-targeting antibody-oligonucleotide conjugates, which facilitate transport of oligonucleotides across the blood-brain barrier (45,46). While these systemic platforms offer a clear administration advantage over lipophilic conjugates and other distribution-enhancing agents requiring direct CNS access, they will also come with new risks, and intrathecal administration remains a clinically viable and highly validated route. When paired with wide-distributing chemistries—such as the D-siRNA discussed here—intrathecal delivery provides a proven path to the clinic that maximizes CNS exposure while minimizing systemic off-target risks.

In conclusion, this work showcases the applicability of the albumin-binding dendrimer conjugate as a safe and effective delivery tool for siRNAs in the CNS. Collectively, the data support safe, wide distribution and efficacious delivery throughout the brain in spinal cord, offering an alternative delivery technology for gene silencing in the brain for the treatment of neurological diseases.

## CONTRIBUTIONS

HHF, MO, JW and AK conceived the project. HHF, MO, JW and AK contributed to the experimental design. HHF MO AS SLS KK RG BMB BM contributed experimentally. HHF, RG, BMB and BM synthesized all siRNA compounds. HHF, MO and JW co-wrote the manuscript. All authors provided feedback and approved of the manuscript.

## CORRESPONDING AUTHORS

Anastasia Khvorova and Jonathan K. Watts.

## DATA AVAILABILITY

All data are presented in the main text and the supplementary information, with raw data available from the corresponding authors upon request.

## ACKNOWLEDGEMENTS

We thank Khvorova/Watts lab members (past and present) for insightful discussions and support. We thank Hanadi F. Sleiman for the initial collaboration on dendrimer conjugate for siRNA delivery, and the productive insights and discussions. We thank the animal medicine core at UMass Chan Medical School, for their help and support on multiple studies. All figures were created with PowerPoint and BioRender.com.

## SUPPLEMENTARY INFORMATION

Supplementary Figures S1– 5 and Supplementary Tables S1.

