## Supplementary Material for "Albumin-binding dendrimer-conjugated siRNA enables safe and effective gene silencing throughout the central nervous system"

**Supplemental Data**

**Supplemental Table 1. Detailed sequences and chemical patterns of oligonucleotides used in this study.** Notations are as follows: "#" –phosphorothioate bond, "m" – 2'-*O*-methyl, "f" – 2'-fluoro, "V" – 5' vinylphosphonate, "Cy3" – Cyanine 3, "Unc" –unconjugated, "Di" – Divalent, "D" – Dendrimer.

| **Gene** | **Oligo ID** | **Sequence (5'-3')** |
|---|---|---|
| *HTT/Htt* | HTT $^{10150}$- Antisense | V(mU)#(fU)#(mA)(fA)(mU)(fC)(mU)(fC)(mU)(fU)(mU)(fA)(mC)(fU)(mG)(fA)(mU)#(fA)#(mU)#(mxU)#(fxU) |
| | HTT $^{10150}$- Sense-Unc | (mU)#(fC)#(mA)(fG)(mU)(fA)(mA)(fA)(mG)(fA)(mG)(fA)(mU)(fU)#(mA)#(fA)(dT)(dT) |
| | HTT $^{10150}$- Sense-Di | (mU)#(fC)#(mA)(fG)(mU)(fA)(mA)(fA)(mG)(fA)(mG)(fA)(mU)(fU)#(mA)#(fA)(dT)(dT)-Divalent |
| | HTT $^{10150}$- Sense-D | (C12)(SB)(C6)(SB)(dT)(dT)(mU)#(fC)#(mA)(fG)(mU)(fA)(mA)(fA)(mG)(fA)(mG)(fA)(mU)(fU)#(mA)#(fA) |
| *JAK1/Jak1* | JAK1$^{860}$- Antisense | V(mU)#(fA)#(mU)(fC)(fG)(fC)(mU)(fU)(mG)(fU)(mA)(fG)(mC)(fU)#(mG)#(fA)#(mU)#(mG)#(mU)#(fC)#(mU) |
| | JAK1$^{860}$- Sense-Unc | CyMN3#(mU)#(mC)#(mA)(fG)(mC)(fU)(mA)(fC)(mA)(fA)(mG)(mC)(mG)(fA)#(mU)#(mA)(dT)(dT) |
| | JAK1$^{860}$- Sense-Di | CyMN3#(mU)#(mC)#(mA)(fG)(mC)(fU)(mA)(fC)(mA)(fA)(mG)(mC)(mG)(fA)#(mU)#(mA)(dT)(dT)-Di |
| | JAK1$^{860}$- Sense-D | (C12)(SB)(C6)(SB)(dT)(dT)(Cy3)#(mU)#(mC)#(mA)(fG)(mC)(fU)(mA)(fC)(mA)(fA)(mG)(mC)(mG)(fA)#(mU)#(mA) |
| *MECP2/Mecp2* | MECP2$^{1764}$- Antisense | V(mU)#(fA)#(mU)(fC)(fG)(fG)(mG)(fA)(mA)(fG)(mC)(fU)(mU)(fU)#(mG)#(fU)#(mC)#(mA)#(mG)#(fU)#(mU) |
| | MECP2$^{1764}$- Sense-Unc | (mA)#(mC)#(mA)(fA)(mA)(fG)(mC)(fU)(mU)(fC)(mC)(mC)(mG)(fA)#(mU)#(mA)(dT)(dT) |
| | MECP2$^{1764}$- Sense-Di | (mA)#(mC)#(mA)(fA)(mA)(fG)(mC)(fU)(mU)(fC)(mC)(mC)(mG)(fA)#(mU)#(mA)(dT)(dT)-Di |
| | MECP2$^{1764}$- Sense-D | (C12)(SB)(C6)(SB)(dT)(dT)((mA)#(mC)#(mA)(fA)(mA)(fG)(mC)(fU)(mU)(fC)(mC)(mC)(mG)(fA)#(mU)#(mA) |
| *APP/App* | APP$^{3265}$- Antisense | V(mU)#(fG)#(mA)(fG)(fA)(fA)(mU)(fU)(mC)(fU)(mU)(fG)(mG)(fU)#(mA)#(fA)#(mU)#(mU)#(mG)#(fA)#(mU) |
| | APP$^{3265}$- Sense-Unc | (mU)#(mU)#(mA)(fC)(mC)(fA)(mA)(fG)(mA)(fA)(mU)(mU)(mC)(fU)#(mC)#(mA)(dT)(dT) |
| | APP$^{3265}$- Sense-Di | (mU)#(mU)#(mA)(fC)(mC)(fA)(mA)(fG)(mA)(fA)(mU)(mU)(mC)(fU)#(mC)#(mA)(dT)(dT)-Di |
| | APP$^{3265}$- Sense-D | (C12)(SB)(C6)(SB)(dT)(dT)(mU)#(mU)#(mA)(fC)(mC)(fA)(mA)(fG)(mA)(fA)(mU)(mU)(mC)(fU)#(mC)#(mA) |
| NTC | NTC Antisense | V(mU)#(fA)#(mA)(fU)(fC)(fG)(mU)(fA)(mU)(fU)(mU)(fG)(mU)(fC)#(mA)#(fA)#(mU)#(mC)#(mA)#(fU)#(mU) |
| | NTC Sense – D | (C12)(SB)(C6)(SB)(dT)(dT)(mU)#(mU)#(mG)(fA)(mC)(fA)(mA)(fA)(mU)(fA)(mC)(mG)(mA)(fU)#(mU)#(mA) |
| | NTC Sense – Unc | (mU)#(mU)#(mG)(fA)(mC)(fA)(mA)(fA)(mU)(fA)(mC)(mG)(mA)(fU)#(mU)#(mA)(dT)(dT) |

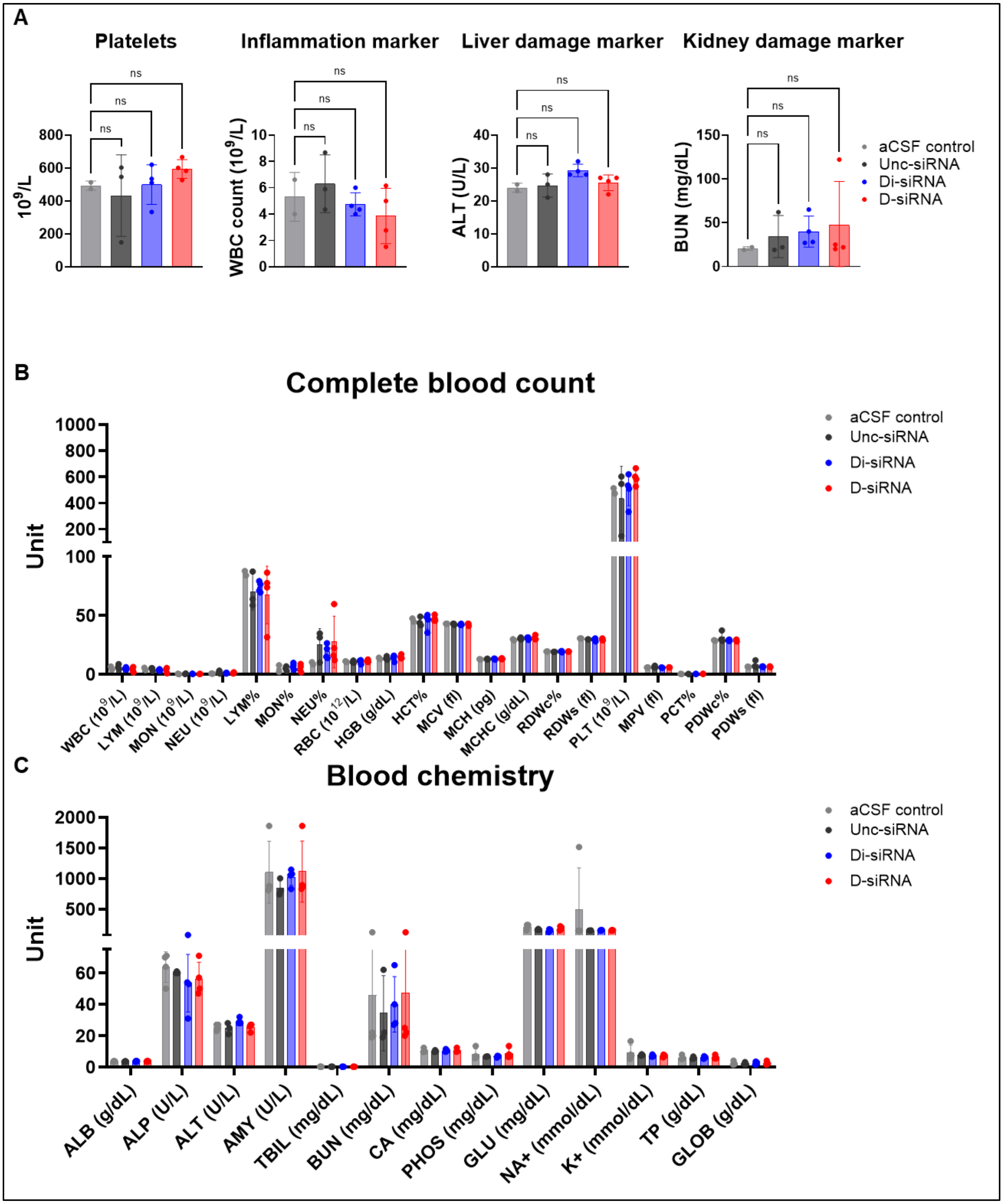


**Supplemental Figure S1. Complete blood count (CBC) and blood chemistry readouts show a favorable safety profile for D-siRNA following CSF administration.** (A) Representative markers from CBC and blood chemistry at 24h post ICV injection, showing no significant changes between the various siRNA chemistries: Platelet count, white blood cells count as marker of inflammation, ALT as marker of liver damage, and BUN as a measure of kidney damage. Full data sets of (B) CBC and (C) blood chemistry readouts. Data represented as mean ± S.D.; n=2-3 mice per group); Statistical analysis was performed in GraphPad Prism: one-way ANOVA analysis of all groups against aCSF control group (ns = non-significant, * p<0.05, ** p<0.01, *** p<0.001, **** p<0.0001).

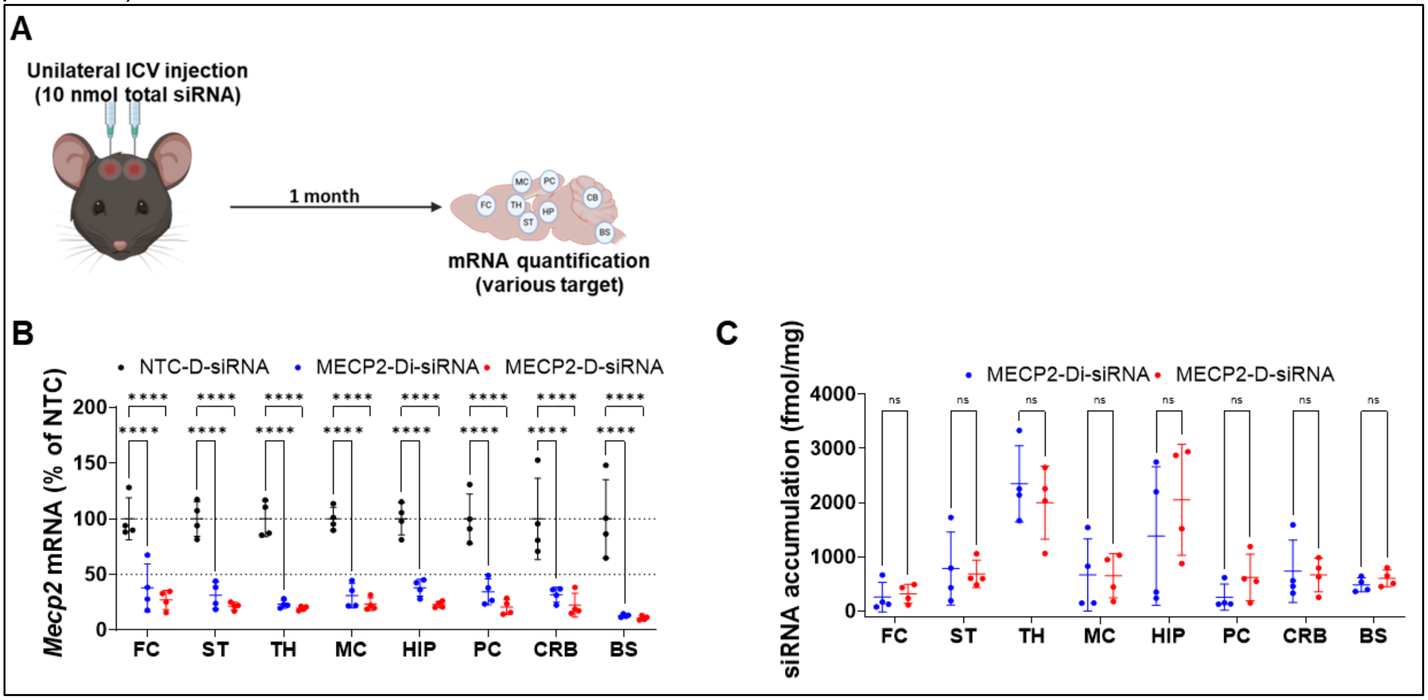


**Supplementary Figure S2. D-siRNA achieves similar efficacy to Di-siRNA targeting *Mecp2* at one-month post ICV injection.** (A) Schematic of *Mecp2*-targeting siRNA efficacy study following bilateral ICV injection of 10 nmol dose at one-month post injection. B) mRNA levels of *Mecp2* compared to NTC-D control and (C) siRNA accumulation levels comparing D-siRNA and Di-siRNA. All study groups are n=4 mice/group. Statistical analysis was performed in GraphPad Prism: two-way ANOVA analysis followed by Dunnet's multiple comparisons of all groups against NTC control group (ns = non-significant, * p<0.05, ** p<0.01, *** p<0.001, **** p<0.0001). (Data represented as mean ± S.D.; mRNA measured by the Quantigene 2.0 Assay; tissue accumulation measured by PNA assay; dotted line on all graphs represents 100% and 50% remaining mRNA protein expression; graphical elements created with Biorender).

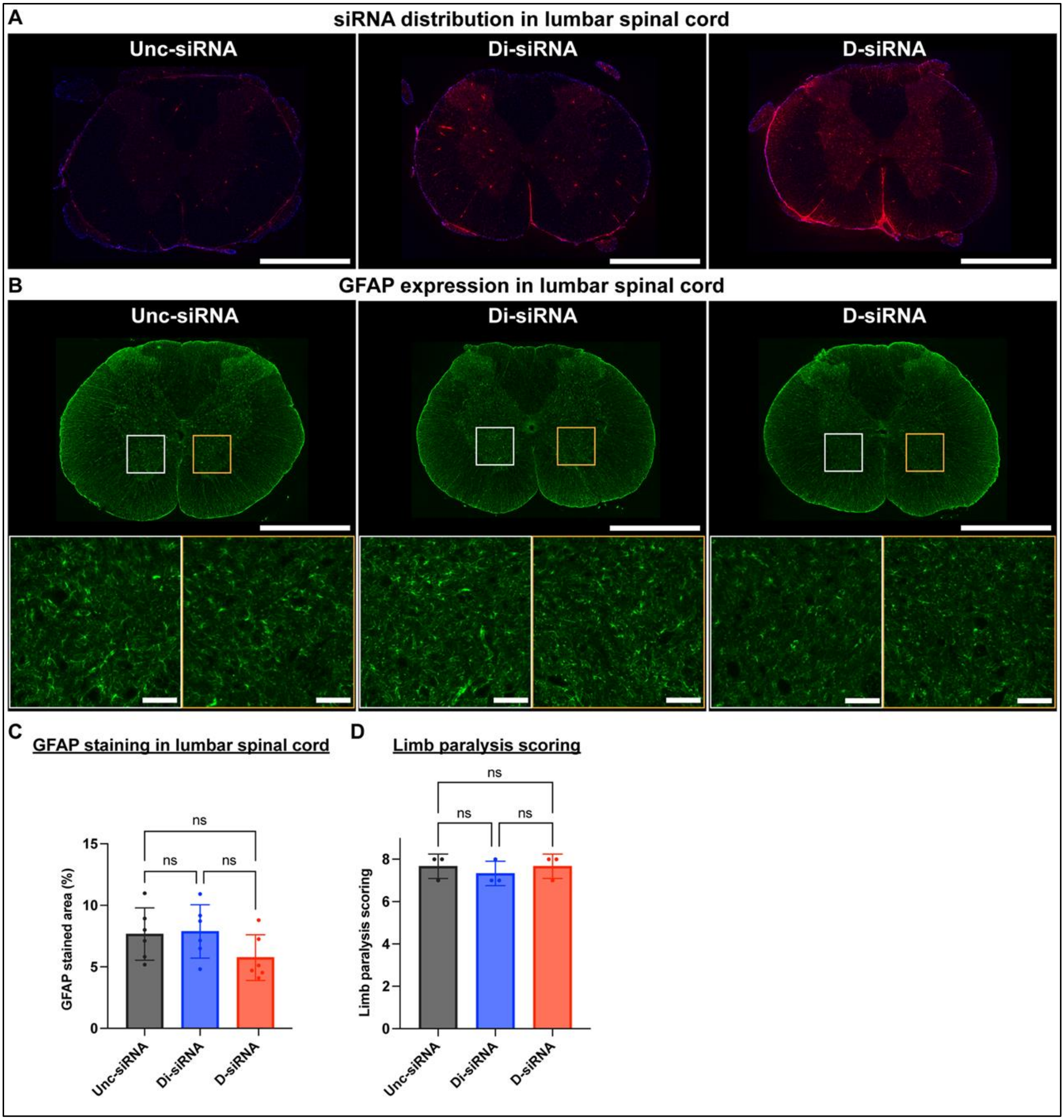


**Supplementary Figure S3. Distribution of siRNA and assessment of astrogliosis in the lumbar spinal cord. (A)** Representative images of the siRNA distribution in the lumbar spinal cord 48 hours after IT injection. Scale bars: 1mm. **(B)** Representative images of GFAP expression in the lumbar spinal cord 48 hours after IT injection. Scale bars: 1 mm (top) and 100 µm (bottom). **(C)** The comparison of GFAP staining area in the lumbar spinal cord. **(E)** The comparison of the limb paralysis scoring 24 hours after IT injection. (Data represented as mean ± S.D.; GFAP quantification by immunostaining, limb paralysis scoring was blinded

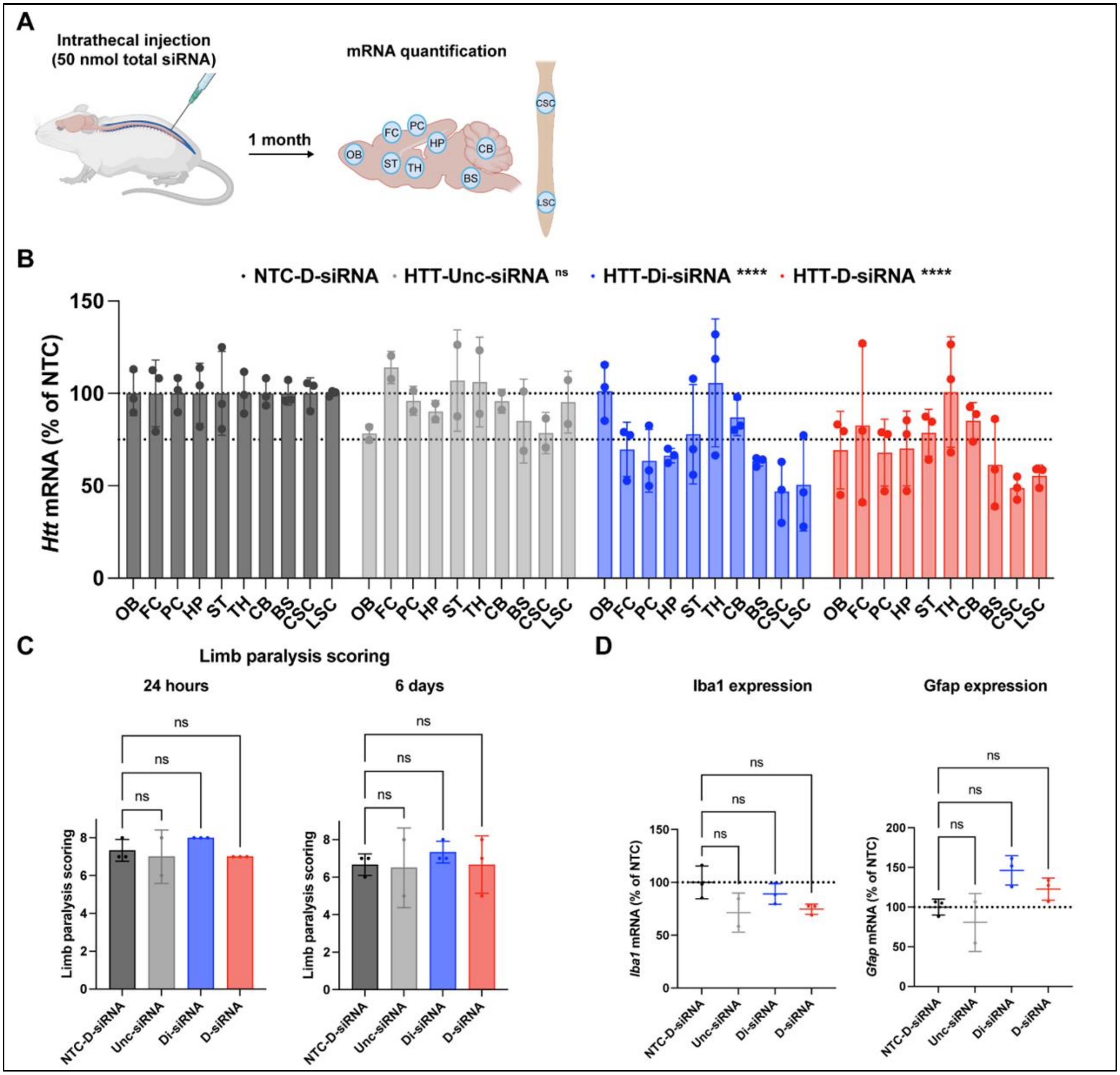


**Supplementary Figure S4. D-siRNA shows comparable silencing of Htt mRNA to Di-siRNA. (A)** Experimental schematic for the study of intrathecal injection of 50 nmol dose of App-targeting Cy3-siRNAs; mRNA levels were assessed 1 month after IT injection of Unc-, Di-, and D-siRNA. **(B)** Silencing data of *Htt* mRNA in various CNS regions. **(C)** Limb paralysis scores at 24 hours and 6 days post-injection. **(D)** Levels of neurotoxicity mRNA markers in the lumbar spinal cord (*Iba1* for microgliosis and *Gfap* for astrogliosis). All study groups are n=2-3 rats/group. Statistical analysis was performed in GraphPad Prism: two-way ANOVA analysis of all groups against NTC control group (ns = non-significant, * p<0.05, ** p<0.01, *** p<0.001, **** p<0.0001). (Data represented as mean ± S.D.; mRNA measured by the Quantigene 2.0 Assay;; dotted line on all graphs represents 100% and 75% remaining mRNA protein expression; graphical elements created with Biorender).

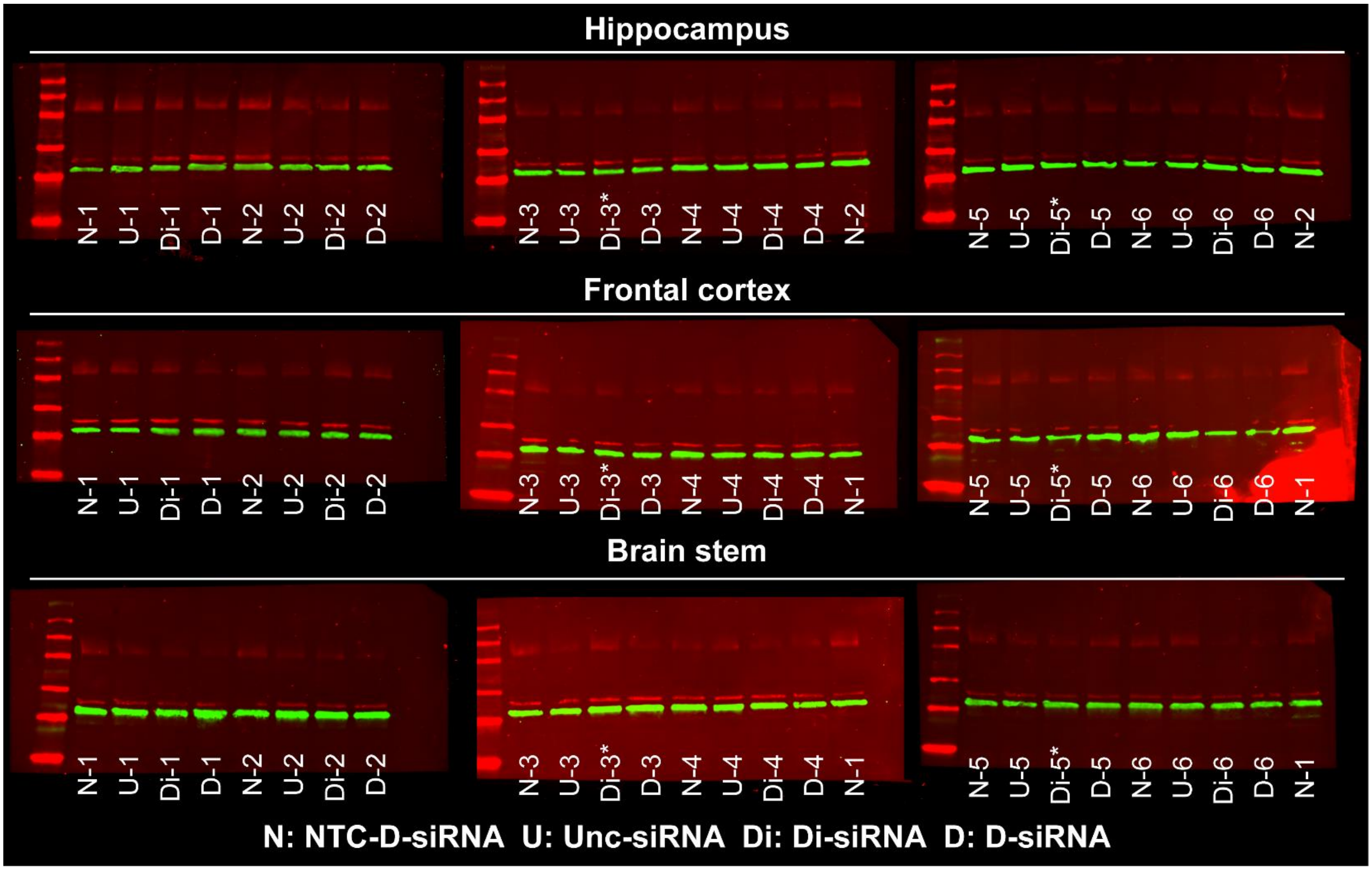


**Supplementary Figure S5.** Original western blot data for Figure 4E.
